# Physicochemical properties of the vacuolar membrane and cellular factors determine formation of vacuolar invaginations

**DOI:** 10.1101/2023.01.17.524399

**Authors:** Yoko Kimura, Takuma Tsuji, Yosuke Shimizu, Yuki Watanabe, Masafumi Kimura, Toyoshi Fujimoto, Miyuki Higuchi

## Abstract

Vacuoles change their morphology in response to stress. In yeast exposed to chronically high temperatures, vacuolar membranes get deformed and invaginations are formed. We show that phase-separation of vacuolar membrane occurred after heat stress leading to the formation of the invagination. In addition, Hfl1, a vacuolar membrane-localized Atg8-binding protein, was found to suppress the excess vacuolar invaginations after heat stress. At that time, Hfl1 formed foci at the neck of the invaginations in wild-type cells, whereas it was efficiently degraded in the vacuole in the *atg8*Δ mutant. Genetic analysis showed that the endosomal sorting complex required for transport (ESCRT) machinery was necessary to form the invaginations irrespective of Atg8 or Hfl1. In contrast, a combined mutation with the vacuole BAR domain protein Ivy1 led to vacuoles in *hfl1*Δ*ivy1*Δ and *atg8*Δ*ivy1*Δ mutants having constitutively invaginated structures; moreover, these mutants showed stress-sensitive phenotypes. Our findings suggest that vacuolar invaginations result from the combination of changes in the physiochemical properties of the vacuolar membrane and other cellular factors.

**Summary statement:** Vacuolar invaginations occur through a combination of the changes of physiochemical properties of the vacuolar membrane and other cellular factors in yeast. Particularly, Hfl1 suppresses the excess invaginations.

## Introduction

Due to the rapid progress of global warming, heat stress on organisms is likely to occur more frequently in the future unless effective countermeasures are taken. When the heat stress is given to cells, many events are initiated or enhanced in a cell or in an organism to decrease the toxic effects and/or to adjust to the high temperature conditions (Morano et al., 2012; Parsell and Lindquist, 1993). For example, protein quality control systems such as molecular chaperones and the protein degradation machinery get enhanced. Moreover, in response to acute and severe heat stress, stress granules, which are composed of translational machinery and RNAs, aim to stop translation tentatively (Protter and Parker, 2016). In addition, heat stress also activates oxidative stress defenses, changes in transport systems, and membrane fluidity . In yeast, cell wall stress pathways are activated as well (Levin, 2011).

In response to chronic and sub-lethal heat stress, yeast acquires thermotolerance over a certain time period (Finley et al., 1987). Several proteins such as ubiquitins and factors involved in the endosomal sorting complex required for transport (ESCRT) are required for survival under chronic heat stress; possibly the cellular systems may be remodeled or reconstructed using ubiquitins and the ESCRT machinery to adapt to such heat stress (Finley et al., 1987; Ishii et al., 2018). However, the precise molecular changes occurring in this state remained unclear.

The vacuole is a single membrane acidic organelle necessary for degrading macromolecules, as well as for nutrient and ion storage, pH homeostasis, and detoxification (Li and Kane, 2009). Moreover, the vacuole is a highly dynamic organelle whose morphology changes in response to various stimuli and situations. In yeast, when cells shift to stationary and starvation phases, as well as in response to hypotonic conditions, the vacuoles fuse. Conversely, in response to hypertonic conditions, they fragment into smaller ones. In addition to fission and fusion, other morphological changes of vacuoles can affect other cell functions (Zhang et al., 2014). For example, in plants, stomatal closure or opening in plants appears to be governed by vacuolar morphological changes in its guard cells (Tanaka et al., 2007). During stomata closing, the vacuole(s) acquire convoluted but continuous structures with decreasing vacuole volume, leading to lower cell volume whereas during stomata opening, the vacuoles acquire less complicated structures with increasing vacuole volume, leading to bigger cell volume.

During heat stress, the vacuoles get deformed and produce invaginations (Numrich et al., 2015b; Ishii et al., 2018). During chronic heat stress, over time the number of vacuoles with invaginations increases and large invaginations often acquire a shape resembling connected vesicles (Ishii et al., 2018). The formation of these invaginations requires ESCRT factors, the SNARE protein Pep2, and ubiquitins. Under heat stress, several plasma membrane proteins are delivered to the vacuole and rapidly degraded by endocytosis (Zhao et al., 2013); under such conditions, more multivesicular bodies (MVBs) are likely delivered and fused with vacuoles. These events would likely lead to formation of vacuolar invaginations. Moreover, we considered that the vacuolar invaginations might represent a cellular strategy for coping with increased vacuolar membranes without increasing vacuole and cell volume.

In addition, we reported previously that formation of vacuolar invaginations is a regulated process (Ishii et al., 2019). Atg8, a member of the ubiquitin-like family and one of the core elements involved in autophagy, suppresses vacuolar invaginations after heat stress (review in (Mizushima, 2020), (Ishii et al., 2019)). The functions of Atg8 could be divided into two categories, autophagy and autophagy-independent functions. Atg8 can be conjugated to lipid phosphatidylethanolamine (PE) through a ubiquitin-like conjugative reaction to form an Atg8–PE complex anchored to the membrane, and the lipidation of Atg8 is necessary for autophagy (Ichimura et al., 2000). Atg8’s autophagy-independent functions, including those in vesicular transport, resistance to oxidative stress, vacuolar fusion, and the formation of lipid bodies, have happened not to require the lipidation of Atg8 (Legesse-Miller et al., 2000; Maeda et al., 2017; Mikawa et al., 2010; Tamura et al., 2010). Since the Atg8’s suppressing function of vacuolar invagination after heat stress does not need its lipidation, it was proposed to be one of autophagy-independent functions (Ishii et al., 2019).

Hfl1 was isolated as an Atg8-binding protein in yeast *Schizosaccharomuces pombe* (Liu et al., 2018). Hfl1 is predicted to be a multi-pass transmembrane protein with seven transmembrane helices and a C-terminal cytosolic tail for Atg8 binding, and it localizes at the vacuolar membrane. Although the *hfl1*Δ mutant does not show autophagy defects, the mutant shows sensitivities against several metal ions including ZnCl_2_, CoCl_2_, and MnCl_2_, and these metal sensitivities shared with the defects of the *atg8*Δ mutant in which its lipidation is unnecessary (Liu et al., 2018; Wang et al., 2014). In another yeast *Sacchromyces cerevisiae*, luminal structures accumulate in the vacuole in the stationary phase in both *atg8*Δ and *hfl1*Δ mutants (He et al., 2021). Thus, Hfl1 has been recognized as a receptor for Atg8 and responsible for the Atg8’s lipidation-independent functions.

In artificial multicomponent giant vesicles (GVs) which are composed of saturated phospholipids, unsaturated phospholipids, and cholesterol, vesicle membrane segregation occurs and domains are created below a miscibility transition temperature (Baumgart et al., 2003; Veatch and Keller, 2005). The domains created by the phase separations of the membrane lipid components have the properties of the liquid-ordered (Lo) or liquid-disorded (Ld) phase. These vesicles further transform their shapes, depending on conditions such as budding (Baumgart et al., 2003; Yanagisawa et al., 2008)

With respect to yeast vacuoles, Lo lipid domains are created in the stationary phase (Rayermann et al., 2017; Toulmay and Prinz, 2013; review in Tsuji and Fujimoto, 2018)’. Lo domains have been shown to mediate microlipophagy (Tsuji et al., 2017; Wang et al., 2014). Moreover, localization analyses of Vph1-GFP, a GFP-tagged integral subunit of the vacuolar-type H^+^ATPase, suggest that the Lo domains, which appear to correspond to Vph1-deficient membrane areas, might be created in response to several stresses, including nutrient deprivation, translation inhibition, weak acids, ER stress, and heat stress (Toulmay and Prinz, 2013; Numrich et al., 2015; Ishii et al., 2018; Liao et al., 2021). In addition, vesicular trafficking to vacuoles plays a critical role in domain formation on the vacuolar membrane (Toulmay and Prinz, 2013). In *S. cerevisiae atg8*Δ and *hfl1*Δ mutants, micro-domain formation as indicated by Vph1-GFP localization has been reported to be defective in the stationary phase (Liu et al., 2018; Wang et al., 2014).

In this study, beginning from the finding that Vph1-GFP distributions in heat-stressed cells are uneven after heat stress, we characterized vacuolar membranes and their invaginations in heat-stressed cells. In addition, we investigated how Hfl1 was involved in vacuolar invagination formation. Our study shows that vacuolar membranes are phase-separated after chronic heat stress and that Hfl1 is localized at the neck of invaginations and involved in suppressing vacuolar invaginations.

## Results

### Phase separation of vacuolar membrane domain formations after heat stress and its correlation with invaginations

In the stationary phase, micro-meter scale domains are created by phase separation of vacuolar membranes, and Vph1-GFP positive area on the vacuolar membrane are stained by Fast Dil which has a higher affinity for a Ld than Lo phase. (Rayermann et al., 2017; Toulmay and Prinz, 2013). A similar Vph1-GFP distribution was observed when shifting log-phase growing cells to higher temperatures (Numrich et al., 2015; Ishii et al., 2018). Besides Vph1-GFP, 12 other vacuolar membrane proteins such as Zrc1-GFP or Ybt1-GFP, show similar distributions in the stationary phase (Toulmay and Prinz, 2013). Similarly, we observed that Zrc1-GFP and Ybt1-GFP showed similar patterns with that of Vph1-GFP after growing cells for 2.5 h at 40.5°C (Fig. 1A). Moreover, when the cell membrane of heat-stressed Vph1-GFP-expressing cells was gently disrupted and the lysates were mixed with Fast Dil, we observed that Fast Dil stained regions of vacuole membrane overlapped with Vph1-GFP positive regions (Fig. 1B). In contrast, in vacuoles from cells grown at 25°C, both Vph1-GFP and Fast Dil showed even distributions (Fig. S1). These results suggested a phase-separation of vacuole membranes occurs after chronic heat stress.

**Fig. 1.**
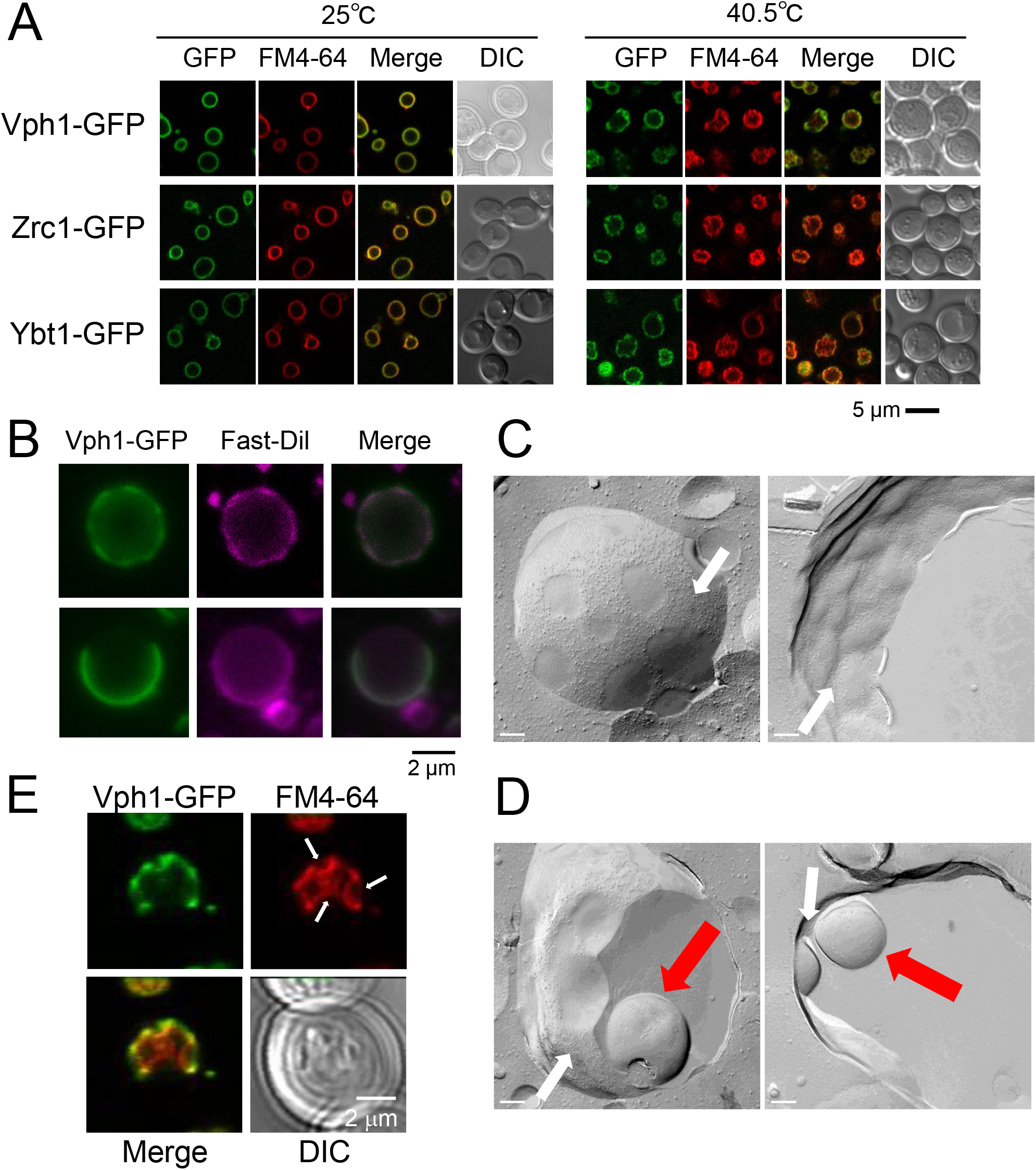
Phase separation, domain formation, and invaginations of vacuole membranes after chronic heat stress at 40.5°C. (A) Localization of Vph1-GFP, Zrc1-GFP, and Ybt1-GFP at 25°C and after 2.5 h of heat stress in wild type cells. GFP and FM4-64 fluorescence of wild-type cells are shown. Scale bar, 5 μm. (B) Phase separation of vacuolar membrane. Two representative images of Fast Dil staining of vacuoles isolated from Vph1-GFP expressing cells after heat stress. Upper and lower panels were vacuoles which were isolated from cells after 2 h and 2.5 h of heat stress, respectively. (C) Freeze fracture EM analysis of vacuolar membrane of wild-type cells after 2.5 h of heat stress, showing that vacuoles membranes segregated into (IMPs)-rich (white arrows) and IMP-less domains. Non-cytoplasmic leaflet (left panel) and cytoplasmic leaflet (right panel) of vacuolar membrane. Scale Bar; 200 nm. (D) Freeze fracture EM analysis of two representative invaginated structures of vacuolar membrane. Vacuolar invaginations and IMP-rich domains are indicated by red and white arrows, respectively. (E) Vacuolar invaginations. FM4-64 staining of 2.5 h heat-stressed Vph1-GFP expressing cells. Vacuolar invaginations are indicated by arrows.

We closely observed the vacuolar membrane in heat-stressed cells by freeze-fracture electron microscopy (EM) (Fig. 1C). After growing them for 2.5 h at 40.5°C, we observed that the vacuole membrane segregated into two different domains, one rich in intramembrane particles (IMPs) and another deficient in IMPs, which looked similar to the vacuolar membrane in the stationary phase. This result supported the above results of the phase separation of vacuolar membrane after heat stress (Moeller et al., 1981; Tsuji et al., 2017). In addition, we observed vacuolar invaginations mainly in IMP-deficient domains (Fig. 1D).

Under heat stress, vacuolar invaginations usually started 2–2.5 h after heat stress, around the same time when uneven Vph1-GFP distribution began to be observed (data not shown). We therefore examined the relationship between vacuolar invaginations and Vph1-GFP localization after heat stress. After 2.5 h of heat stress, at 75% (SE±3.5%) of invagination areas, more Vph1-GFP was localized in the neck region of vacuolar membranes than on the inside of the invagination, suggesting that vacuolar invaginations are mainly induced by phase separation of the vacuolar membrane (Fig. 1E). These results indicated that heat stress induced phase separation of the vacuole membrane in yeast, in turn inducing vacuolar invaginations.

### Increased vacuolar invagination in *hfl1*Δ cells after chronic heat stress

Hfl1, an Atg8 binding protein, has been proposed to mediate the functions of non-lipidated form of Atg8 (Liu et al., 2018). To examine the vacuole morphology of *hfl1*Δ mutants after chronic heat stress, we used cytoplasmic Pgk1-GFP-expressing cells as vacuolar invaginations are easily detected by GFP fluorescence in the cytosol and stained vacuole membranes with FM4-64 (Fig.2). At 25°C, many vacuoles showed normal morphologies, but we observed that a few cells already contained vacuole(s) with invaginations (Fig.2D, Fig. S2). After shifting the temperature to 40.5°C, we observed a drastic change in vacuole morphology: vacuolar invaginations drastically increased as in the *atg8*Δ mutant. These results suggested that both Atg8 and Hfl1 are involved in suppressing excessive vacuolar invaginations after chronic heat stress.

**Fig. 2.**
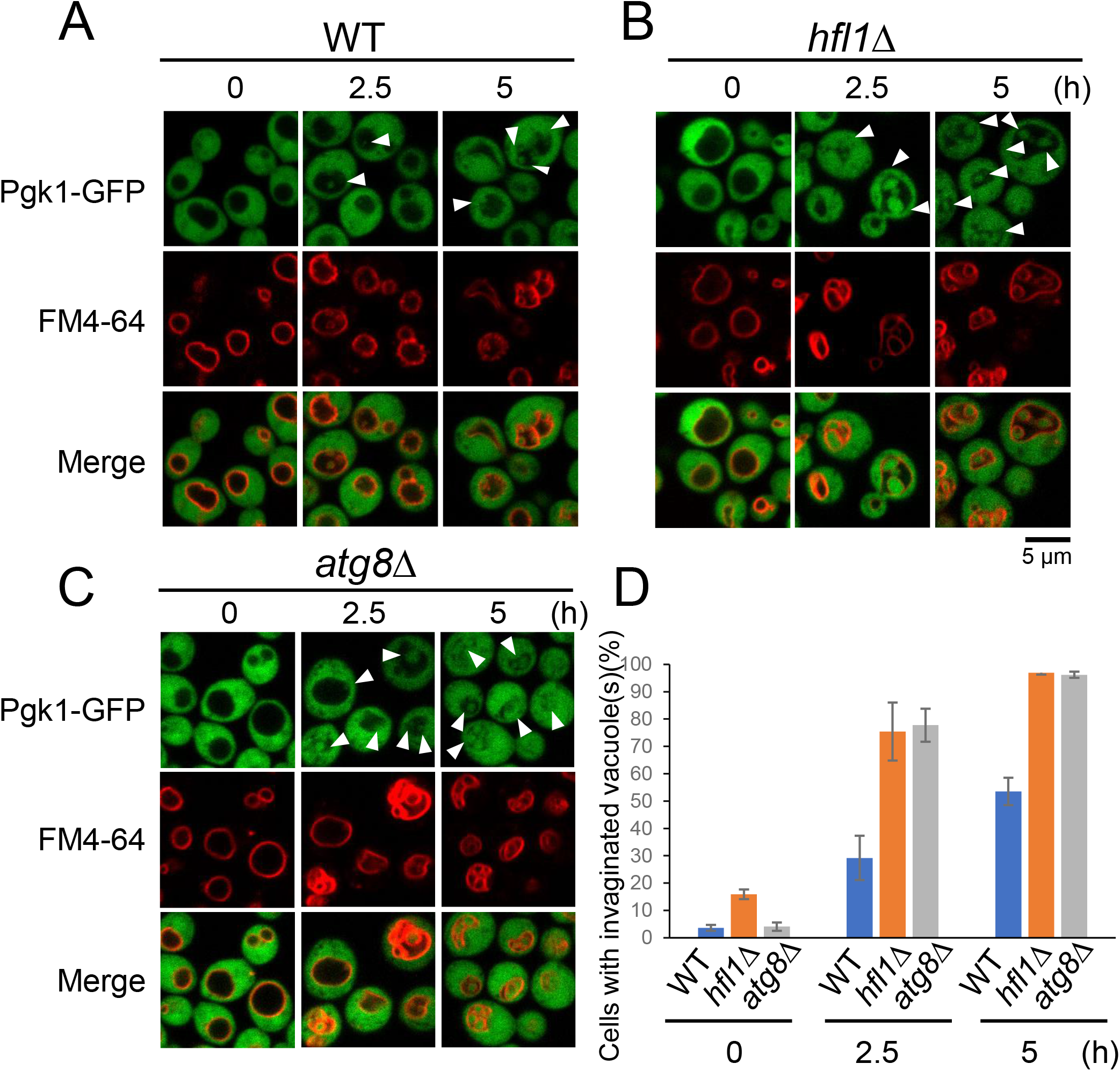
Greater acceleration of invaginations in the vacuolar membranes of *hfl1*Δ after chronic heat stress. GFP and FM4-64 fluorescence of wild-type (A), *hfl1*Δ (B) and *atg8*Δ (C) cells, expressing Pgk1-GFP at 25°C and 40.5°C for 2.5 and 5 h. Scale bar, 5 μm. Vacuolar invaginations are indicated with arrows. Because FM4-64 fluorescence was fainter in cells at 25°C than in cells after heat stress, the contrast of the images of cells at 25°C was enhanced. (D) Quantification of (A), (B) and (C). Cells with invaginated vacuoles were counted. Data indicates mean ± standard error of the mean (SE). Statistical significance: p = 0.028 and 0.011 by both-sided t-test for the pairs between wild-type and *hfl1*Δ cells at 2.5 and 5 h, respectively. No significant difference was observed for the pairs between *hfl1*Δ and *atg8*Δ cells either at 2.5 or 5 h.

We then examined Vph1-GFP localization of *atg8*Δ and *hfl1*Δ mutants after heat stress to investigate whether domain formation occurred on the vacuolar membrane in these mutants (Fig.3A). After 2.5 hr at 40.5°C, some vacuoles showed unevenly distributed Vph1-GFP suggesting formation of domains in both of the mutants, but these uneven Vph1-GFP distributions were not clearly observed in the vacuoles with invaginations, possively due to the accelerated invaginations.

**Fig. 3.**
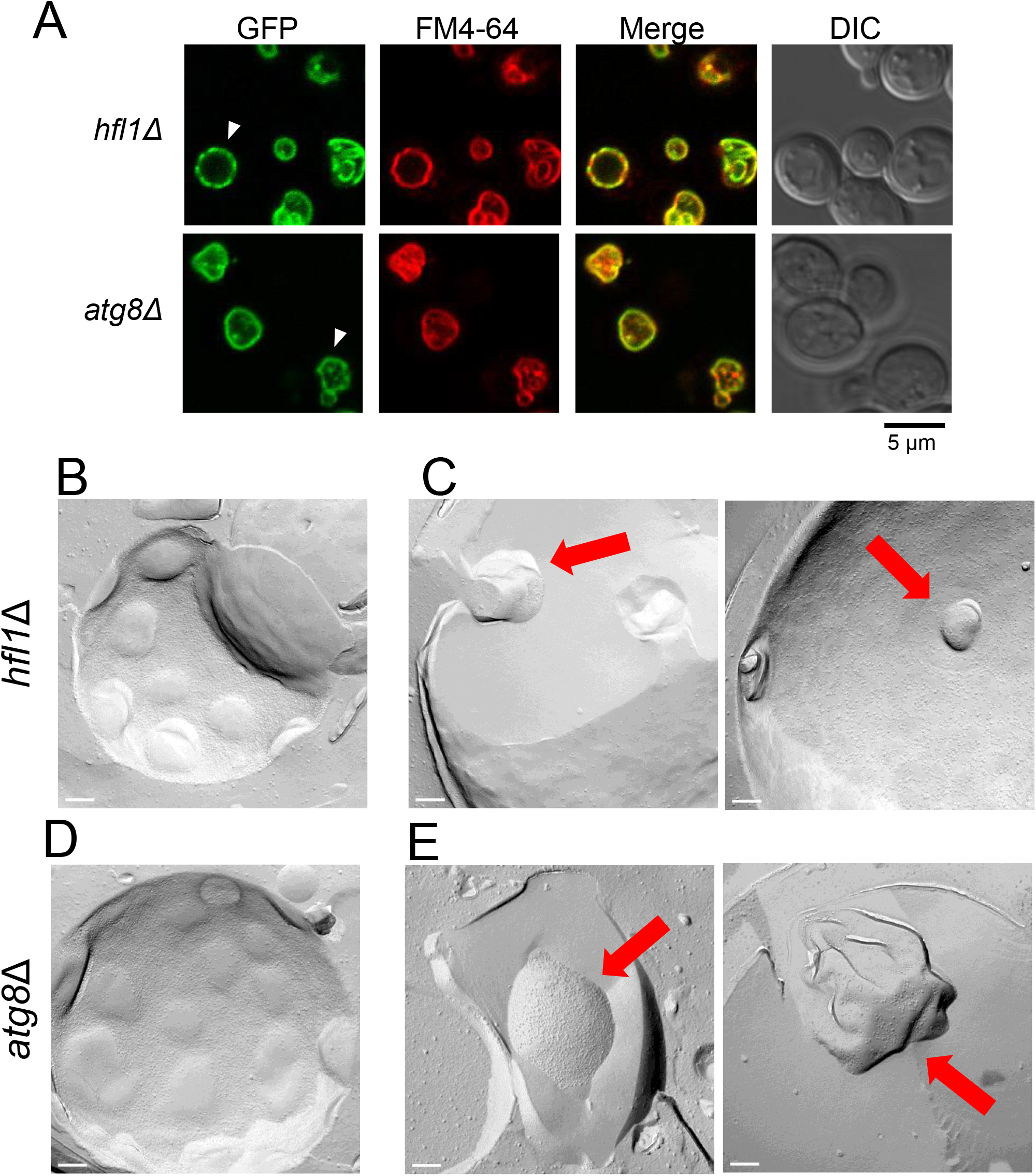
Domain formation of vacuole membranes in *hfl1*Δ and *atg8*Δ cells after chronic heat stress. (A) Localization of Vph1-GFP after 2.5 h of heat stress. Scale bar, 5 μm. Arrowhead indicates a vacuole with uneven Vph1-GFP distribution. (B) Freeze fracture EM analysis of vacuole domains of *hfl1*Δ cells. Scale Bar; 200 nm. (C) Freeze fracture EM analysis of vacuolar invaginations in *hfl1*Δ cells. Vacuolar invaginations are indicated by red arrows. Two images are shown. (D) Freeze fracture EM analysis of vacuole domains in *atg8*Δ cells. (E) Freeze fracture EM analysis of vacuolar invaginations in *atg8*Δ cell. Two images are shown.

Freeze fracture EM analysis was performed to observe the vacuole membrane of these mutants. In both *hfl1*Δ and *atg8*Δ mutants, IMP-deficient domain-like regions like those of wild-type cells were clearly observed after 2.5 h of heat stress (Fig.3B and D). However, vacuolar invaginations occurred irrespective of the vacuole membrane region (Fig.3C and E). In addition, the shape of invaginations was varied and irregular. These results suggest that in *hfl1*Δ and *atg8*Δ mutants, vacuolar domain formations occurred but invaginations did not follow the domains, taking aberrant shapes.

### Localization of Hfl1-ymNeonGreen foci at the neck of the invagination

To understand Hfl1 function on vacuolar invaginations after heat stress, we examined Hfl1 localization. We created a strain in which a fusion of Hfl1 and ymNeon Green (NG), a brighter fluorescence protein than GFP, was endogenously expressed under the *HFL1* promoter. This construct enabled us to detect the Hfl1 localization at endogenous levels (Fig.4). At 25°C, Hfl1-NG was distributed overall on the vacuolar membrane; in about 30% of cells, one Hfl1-NG focus was mostly observed on the vacuole membrane (Fig.4A, B and C). These foci were mainly localized on the smooth vacuolar membrane and some at the junction between two vacuoles (Fig.4D, Fig. S3). At 4 h after heat stress, we observed more cells contained ≥1 Hfl1-ymNG foci than cells grown at 25°C with more foci per vacuole (Fig.4B and C). In addition, we observed >50% foci were localized at the neck of invaginations (Fig.4A and D).

**Fig. 4.**
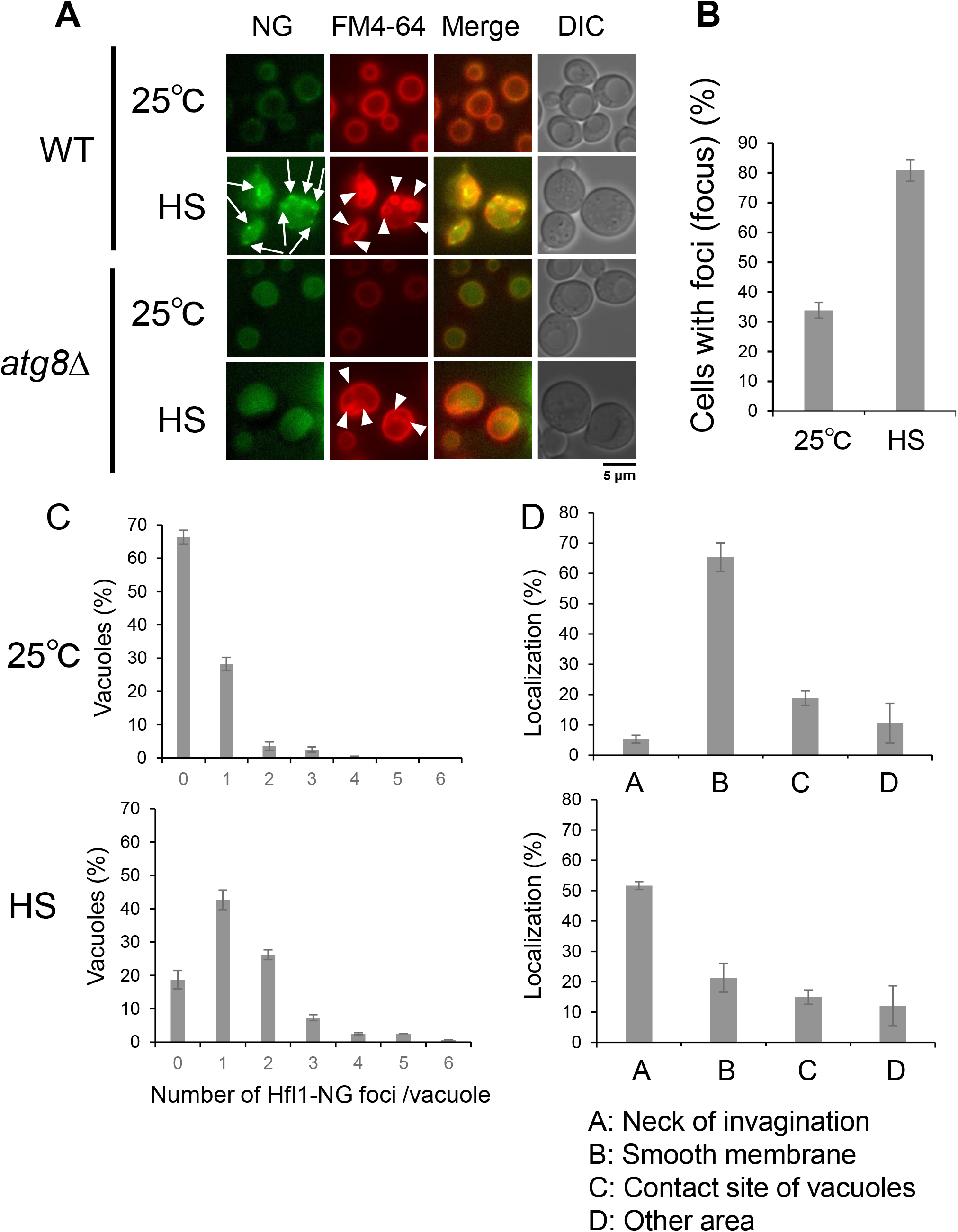
Hfl1 localization. (A) Localization of Hfl1-ymNeonGreen (Hfl1-NG) in wild-type cells and *atg8*Δ cells grown at 25°C and 40.5°C for 4 h. Scale bar, 5 μm. NG and FM4-64 fluorescence, and DIC images. Arrow and arrowhead indicate a Hfl1-NG focus and an invagination, respectively. (B) Ratio of cells harboring vacuoles with Hfl1-NG foci grown at 25°C and 40.5°C for 4 h in wild-type cells. Hfl1-NG foci were counted in > 40 cells per experiment in >3 independent experiments. The mean ± SE is shown. (C) Number of Hfl1-NG foci per vacuole grown at 25°C (upper graph) and 40.5°C for 4 h (lower graph) in wild-type cells. (D) Foci localization of wild-type cells grown at 25°C (upper graph) and 40.5°C for 4 h (lower graph) in wild-type cells. Each focus in three independent experiments was examined for their localization on a vacuole.

In *atg8*Δ mutant grown at 25°C, we observed a weaker NG fluorescence on the vacuolar membrane and the inside of vacuoles, with hardly any foci. After heat stress, the fluorescence was mainly detected inside the vacuole (Fig.4A). These results suggested that Atg8 promoted Hfl1-NG foci formation and that some Hfl1-NG was delivered from the vacuolar membrane to the inside of the vacuole in *atg8*Δ mutants grown at 25°C; a delivery enhanced after heat stress.

### Hfl1 degradation after heat stress in *atg8*Δ mutants

We examined the expression of an endogenous Flag-tagged Hfl1, Hfl1-3xFlag (Fig.5, Fig. S4) after submitting cells to heat stress for 3 h. In these conditions, a massive decrease of Hfl1-3xF levels was observed in *atg8*Δ cells whereas an only small decrease was detected in wild-type cells. This decrease in *atg8*Δ cells was cancelled in *atg8*Δ*pep4*Δ mutant in which a gene encoding Pep4, a major vacuolar protease, was additionally disrupted, indicating that Hfl1 was degraded in the vacuole. Moreover, the levels of an Hfl1-3xFlag mutant with changes to four amino acids (W371A, I375A, D384A, Y387A), responsible for Atg8 binding (Liu et al., 2018), greatly decreased after heat stress even in the presence of Atg8. These results indicated that Atg8 protects Hfl1 from degradation by direct interaction and that without Atg8, Hfl1 is efficiently degraded in the vacuole after heat stress.

**Fig. 5.**
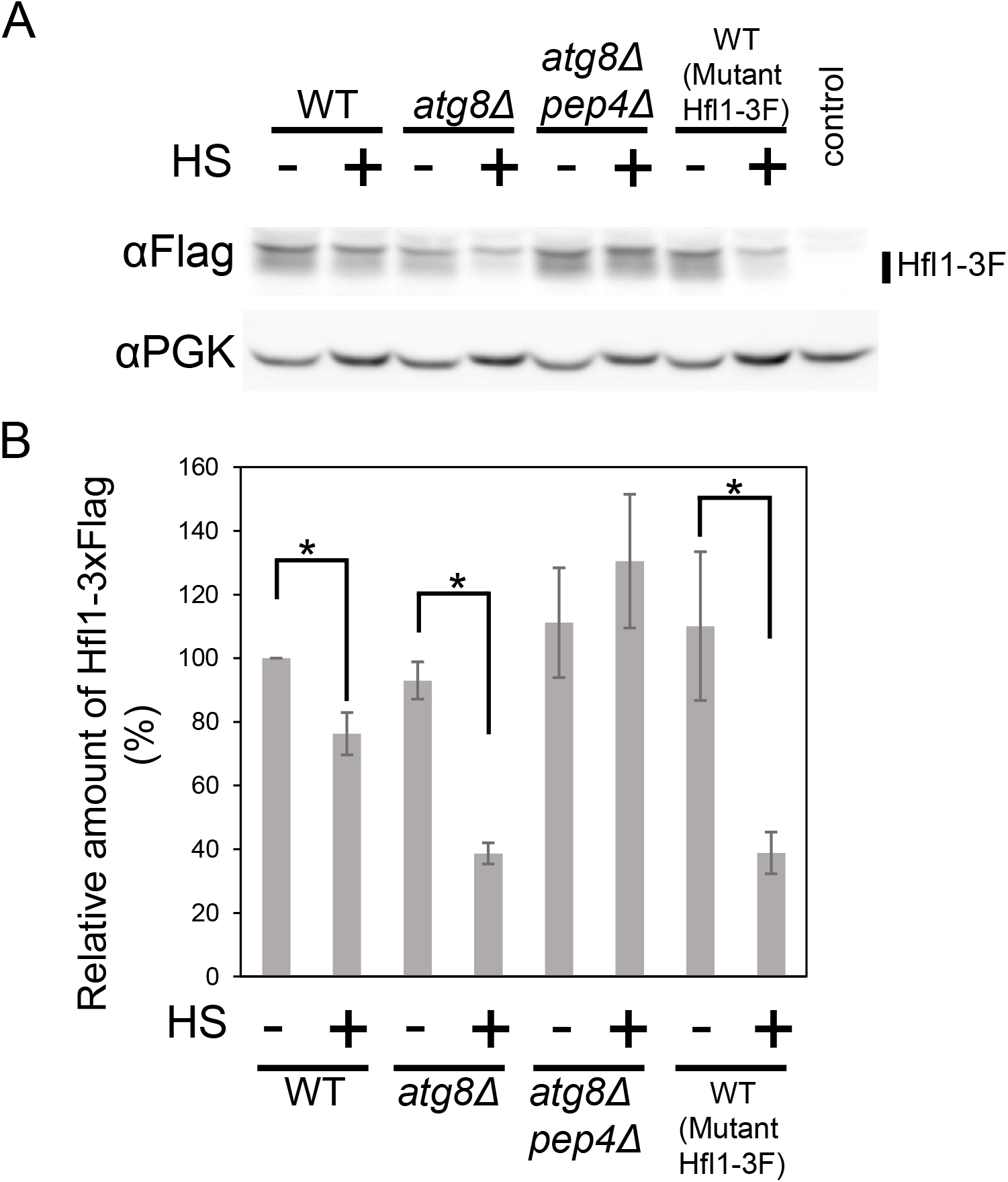
Hfl1-3xFlag expression. (A) Expression of Hfl1-3xFlag and Pgk1 in wild-type, *atg8*Δ, *pep4*Δ *atg8*Δ cells, and cells expressing mutant Hfl1-3xFlag (W371A, I375A, D384A, Y387A) grown at 25°C and 40.5°C for 3 h. A control strain not expressing Hfl1-3xFlag is shown at the rightmost lane. (B) Quantification of (A).

### Relationship between Hfl1 and ESCRTS

Next, to understand Hfl1 functions with respect to other cellular factors, we combined an *hfl1*Δ mutation with other mutations and observed the vacuolar morphology at normal temperature as well as after heat stress.

Vacuolar membrane invaginations are not produced after chronic heat stress in mutants of ESCRT factors such as Vps23 (ESCRT-I) and Vps24 (ESCRT-III), and they are sensitive to chronic heat stress (Ishii et al., 2018). In addition, in *atg8*Δ*vps24*Δ and *atg8*Δ*vps23*Δ double mutants vacuolar invaginations are severely impaired after heat stress, resembling the phenotype of *vps24*Δ and *vps23*Δ, indicating that *vps24*Δ and *vps23*Δ are epistatic to *atg8*Δ. We found that like in these mutants, the vacuolar invaginations of *hfl1*Δ*vps24*Δ were severely impaired after heat stress (Fig.6). Similarly, vacuolar invaginations were defective in *hfl1*Δ*vps23*Δ and *hfl1*Δ*vps4*Δ mutants (Fig. S5). These results suggest that mutations for ESCRT components are epistatic to both *atg8*Δ and *hfl1*Δ.

**Fig. 6.**
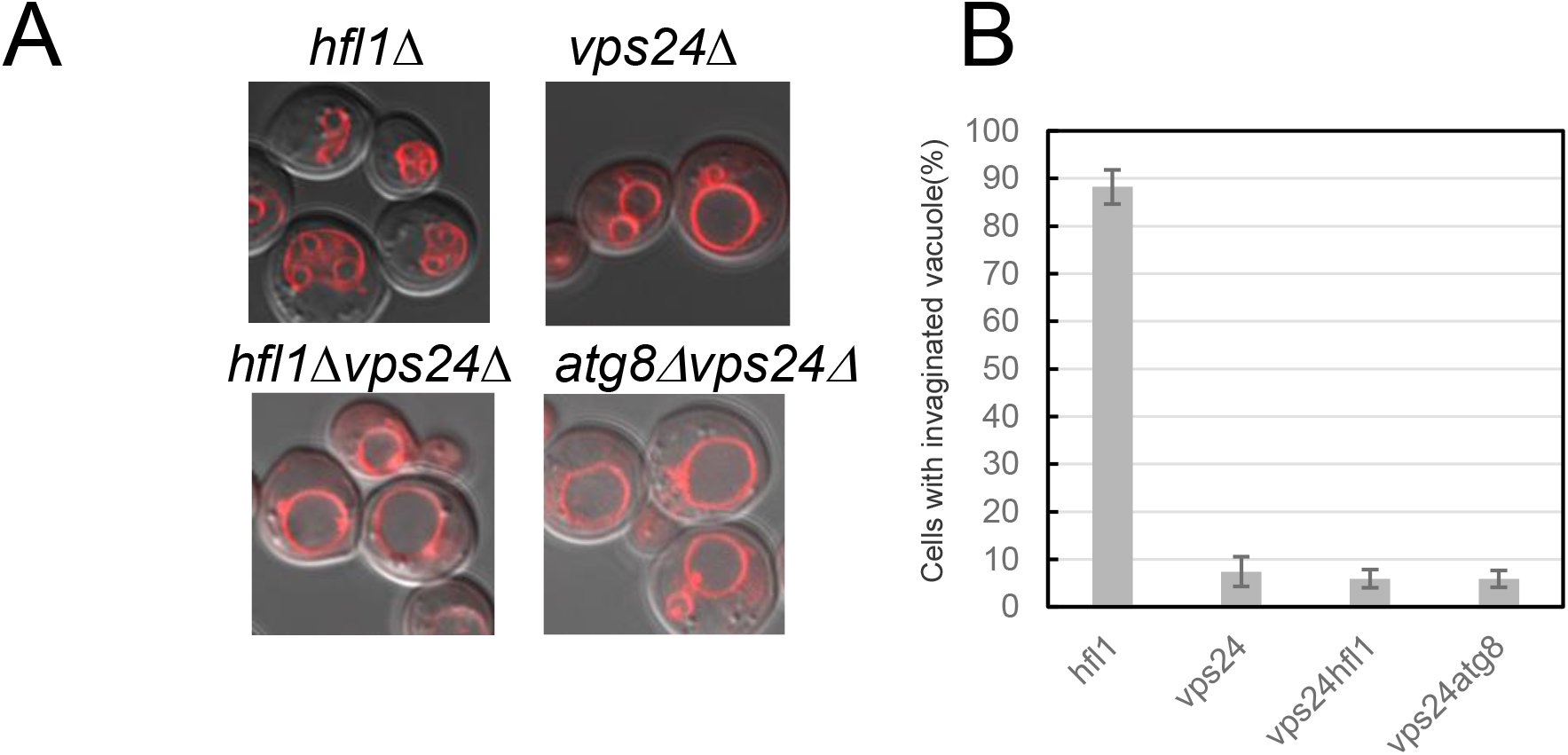
Relationship of *hfl1*Δ with mutants for ESCRT factors. (A) Vacuolar morphologies of wild-type, *vps24*Δ, *hfl1*Δ, *atg8*Δ, *hfl1*Δ*vps24*, and *atg8*Δ*vps24*Δ cells grown at 40.5°C for 4 h. Merged FM4-64 fluorescence and DIC images. (B) Quantification of cells with invaginated vacuoles in (A). Cells with invaginated vacuole structures were counted ≥50 cells in each experiment in three independent experiments. The mean ± SE is shown.

### Constitutive vacuolar membrane invaginations in *hfl1*Δ*ivy1*Δ mutants

Furthermore, we examined whether Hfl1 was related to Ivy1. Ivy1 is a vacuole-localized protein with an Inverted-BAR (I-BAR) domain. An *atg8*Δ*ivy1*Δ double mutant shows vacuoles with constitutive massive invaginations even at 25°C (Ishii et al., 2019). We investigated whether *hfl1*Δ*ivy1*Δ double mutant showed a similar vacuolar morphology. Indeed, we observed highly invaginated vacuolar structures in the *hfl1*Δ*ivy1*Δ mutant (Fig.7A and B). However, Vph1-GFP distributed evenly in these mutants, suggesting absence of domains (Fig.7C).

**Fig. 7.**
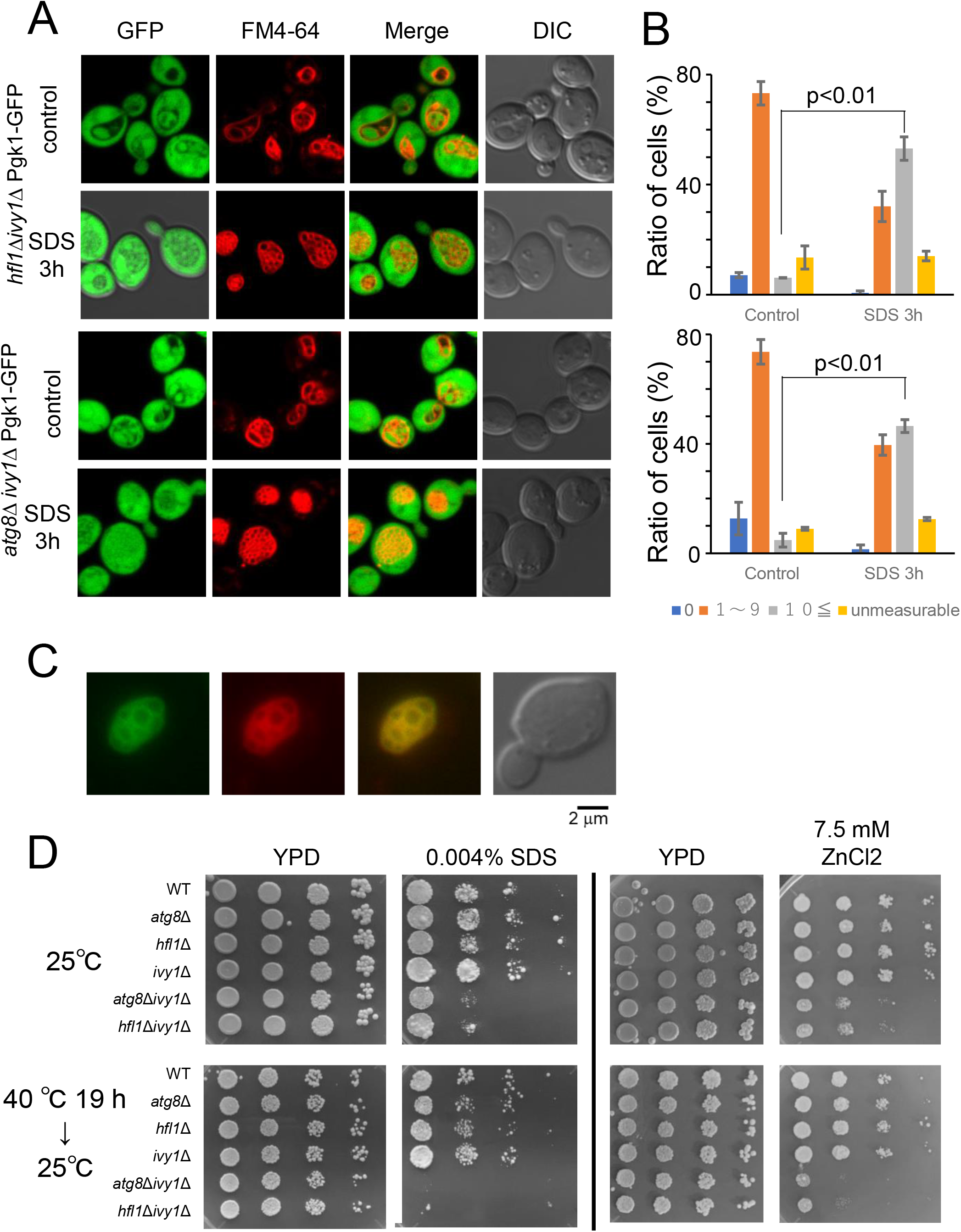
Constitutive vacuolar invaginations and stress sensitivities of *hfl1*Δ*ivy1*Δ cells. (A) Vacuolar morphologies of *hfl1*Δ*ivy1*Δ and *atg8*Δ*ivy1*Δ cells expressing Pgk1-GFP grown at 25°C and after 3 h of SDS treatment at 25°C. Scale bar, 5 μm. (B) Quantification of vacuolar invaginations of *hfl1*Δ*ivy1*Δ and *atg8*Δ*ivy1*Δ cells with or without SDS treatment. The vesicle-like structures inside of vacuoles were counted. The experiment was repeated three times and the mean ± SE is shown. (C) Localization of Vph1-GFP of *hfl1*Δ*ivy1*Δ cells grown at 25°C. GFP and FM4-64 fluorescence, and DIC are shown. Scale bar, 2 μm. (D) Sensitivities against SDS and ZnCl2. Serial 10-fold dilutions of indicated strains grown at log phase were spotted on YPAD, YPAD+0.004% SDS, YPAD+5.0 mM ZnCl2, and YPAD+7.5 mM ZnCl2 plates. Cells were grown at 25°C or at 40°C for 18 h followed by the incubation at 25°C. Cells were incubated at 25°C for 3 days for YPAD, 6 days for SDS-containing plates, and 8 days for ZnCl2 containing plates.

Similar to the *atg8*Δ*ivy1*Δ mutant, *hfl1*Δ*ivy1*Δ mutant growth was reduced at high ZnCl_2_ concentrations, which would affect vacuolar activity (Fig.7D) (MacDiarmid et al., 2003). In addition, *hfl1*Δ*ivy1*Δ and *atg8*Δ*ivy1*Δ mutants exhibited a severe growth defect when plated on a medium containing SDS, which perturbates both the plasma membrane and cell walls, whereas a single mutation, *hfl1*Δ, *ivy1*Δ, or *atg8*Δ, did not cause such sensitivity. These defects were increased when cells were plated on ZnCl_2_ or SDS-containing media and heat-stressed for 18 h before incubation at 25°C. Moreover, we observed that vacuole invaginations became further intricated with SDS treatment in mutants grown at 25°C (Fig.7A).

Collectively, these results suggest that Atg8 or Hfl1 are required in the *ivy1*Δ mutant to suppress vacuolar membrane invaginations at 25°C. Moreover, the maintenance of vacuolar membrane homeostasis, avoiding excess invaginations, appears to be important for cell growth.

## Discussion

### Phase separation and invagination of vacuolar membrane after heat stress

In this study, we showed phase-separation of the vacuolar membrane occur in heat stressed cells, and lipid domains are created on the vacuole membrane. Moreover, since more Vph1-GFP localized at the neck of the invaginated area than at the inside of the invagination, we suggest that the phase-separation of vacuolar membrane induces the invaginations (Fig.8). Biophysical analyses of artificial tertiary-component GVs show that lipid domain formations, caused by phase separation of different lipids, lead to the formation of various shape such as budding. A similar process might occur in the vacuole membrane of heat-stressed cells, the invagination may decrease a line energy of these domain boundaries, caused by phase separation of the membrane. By having such invaginations, the cell would avoid the massive increase of the vacuole volume, which would result in the increase of the cell volume.

**Fig. 8.**
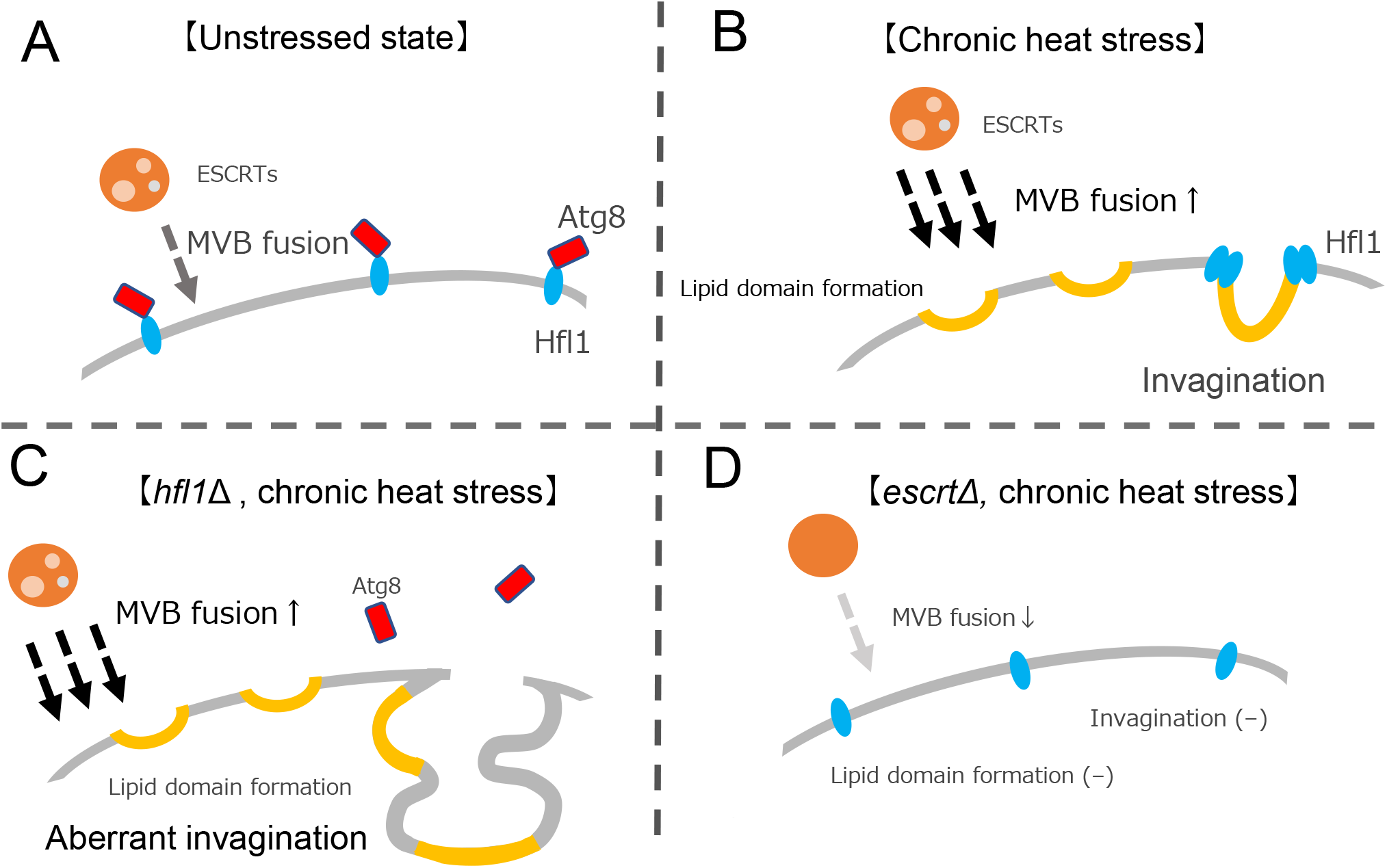
Model of vacuolar membrane invagination and Hfl1 and Atg8 role in response to chronic heat stress. At normal temperatures, Hfl1 is distributed in the vacuole membrane and binds to Atg8 (Panel A). With heat stress, more MVB vacuoles fuse with vacuoles than in the normal state and the vacuole membrane phase-separates forming invaginations according to the phase-separated domains (Panel B). Hfl1 accumulates at the neck of the invaginations and suppresses the excess. However, whether Atg8 is localized at the Hfl1 foci after heat stress is currently unclear. In the *hfl1*Δ mutants, the vacuolar membrane phase-separates but the invaginations occur irrespective of phase-separated domains (Panel C). Similar scenario are expected in the *atg8* Δ mutants, because Hfl1 is degraded after heat stress in the mutant. In ESCRT mutants, MVB formation is impaired, the vacuolar membrane does not phase-separate, and invaginations do not occur after heat stress (Panel D). Hfl1 localization and whether Atg8 is localized with Hfl1 have not been investigated in ESCRT mutants after heat stress.

Although phase separation appears to occur in the vacuole membranes of heat-stressed cells as well as in the stationary phase and GV, there is a difference in the temperatures for phase separation. In this study, we found that phase separation of the vacuolar membrane occurs with increasing temperature whereas for GV and vacuole membrane in the stationary phase, phase separation occurs upon decreasing temperatures (Lipowsky, 1992; Rayermann et al., 2017; Leveille et al., 2022). We speculate that these discrepancies could be explained because in the case of GV and vacuoles from stationary phase cells, there is barely any materials exchange in these membranes and phase separation depends on the physical properties of existing molecules in the membrane. In contrast, yeast cells are in a log phase and active when heat stress is applied. Thus, the vacuolar membrane compositions would change to cause phase-separation. Indeed, ubiquitin-deficient mutant (*ubi4*Δ) and ESCRT mutants show homogeneously distributed Vph1-GFP fluorescence on the vacuolar membranes after heat stress, suggesting the phase separation does not occur in these mutants (Ishii et al., 2018) (Fig.8). During heat stress, fusion between endosome and vacuole would occur more often in heat-stressed cells and materials such as sterols are released from the intraluminal vesicles of MVBs in the vacuole lumen to be delivered to the vacuolar membrane. In addition, homeoviscous adaptation, the production of more saturated lipids and less unsaturated lipids when temperature increases might affect the vacuoles as well (Sinensky, 1974; Fan and Evans, 2015). Altogether, these activities might affect vacuolar membrane composition, resulting in phase-separation.

### The possible function of Hfl1

In contrast with the spontaneous budding or membrane of multicomponent GVs, vacuole invaginations after heat stress appear to respond to a regulated process. We showed that Hfl1 and Atg8 suppressed invagination formation. In addition, Hfl1-NG foci were localized at the neck of the invaginations after heat stress. Since phase-separation seems to occur in heat-stressed vacuolar membranes, Hfl1 may recognize and regulate these lipid states of the vacuolar membrane. The accumulation of Hfl1 at the neck of the invaginations suggest that Hfl1 might exist at the boundary of the Lo and L domains, reducing the line energy of the boundary of Lo and Ld areas of phase-separated membranes, somehow suppressing indiscriminate invaginations.

Hfl1-positive foci may contain other molecules important for suppression of invaginations. Atg8 does not seem to be included in Hfl1 foci as we could not detect GFP-Atg8 foci at the neck of the vacuolar membrane after heat stress (Ishii et al., 2019). However, since Hfl1 is expressed at very low levels, the Hfl1-Atg8 complex might not be detected. Moreover, since we could not precisely capture the moment of vacuolar invagination, it is not clear whether Hfl1 foci localization occurs before or after formation of vacuolar invaginations. Further research is required to elucidate the molecular functions of Hfl1 and the mechanism of Hfl1 foci formation.

Our finding that Hfl1 was degraded in the vacuole of the *atg8*Δ mutant after heat stress, indicates that Atg8 might stabilize Hfl1 protein before forming foci. In addition, since Hfl1-NG foci was hardly detected in the *atg8*Δ mutant, Atg8 might function in Hfl1 foci formation. The GFP fluorescence of overexpressed Hfl1-GFP was reported to be weaker in an *atg8*Δ mutant than in wild-type cells in the stationary phase (He et al., 2021). This suggests that Hfl1 might also be less stable in the absence of Atg8 in the stationary phase. However, the molecular mechanism underlying delivery of Hfl1 to the inside of the vacuole in *atg8*Δ mutants is currently unknown. One possible mechanism would be microautophagy, by which vacuolar membrane proteins are directly internalized into the vacuole; a possibility which should be investigated in the future (Morshed et al., 2020; Oku et al., 2017; Zhu et al., 2017).

### Biological significance of vacuolar invaginations

So far, we did not observe growth defects in *hfl1*Δ and *atg8*Δ mutants after heat stress (data not shown); therefore, it was unclear whether suppressing excess invaginations was important to the cell. However, constitutive vacuolar invaginating mutants of *atg8*Δ*ivy1*Δ and *hfl1*Δ*ivy1*Δ were stress sensitive, suggesting the importance of having adequate vacuolar invaginations.

Interestingly, based on Vph1-GFP patterns, domains did not seem to be created in *hfl1*Δ*ivy1*Δ mutants. Although the reason why these mutants showed these phenotypes is unclear, we consider that irrespective of vacuolar membrane phase-separation, invaginations are regulated by factors such as Atg8 or Hfl1 and Ivy1, which are required to suppress invagination even at normal temperatures and to keep vacuoles in their spherical form. The I-Bar domain in Ivy1 has been reported to bind to the membrane curvature (Salzer et al., 2017). Ivy1 has been reported to be localized at the negative curvature of invagination after heat stress, though Ivy1 is dispensable for vacuolar invagination after heat stress (Numrich et al., 2015a). The vacuoles in *ivy1*Δ*vma16*Δ mutants was reported to have membranous structures at normal temperatures, which suggests that Hfl1, Atg8, and Vma16 may share a common function in suppressing invagination in conjunction with Ivy1 (Numrich et al., 2015).

Finally, as yeast is a eukaryotic model organism, we believe that a similar vacuolar phenomenon might occur in the cells of diverse organisms under stress as well as in normal conditions. Moreover, similar to the structural changes in the vacuoles of guard cells involved in the stomatal movement, structural changes to vacuoles might affect cellular/systemic biological activities. The present results can contribute to the understanding of biological activities associated with changes in vacuolar structures.

## Methods

### Media, yeast strains and plasmids

Yeast strains were grown in YPAD medium [1% yeast extract, 2% Bacto–Peptone or Hipolypepton (Nihon Seiyaku), 2% glucose, and 0.002% adenine], in synthetic complete medium (SD; 0.67% yeast nitrogen base and 2% glucose supplemented with amino acids), or synthetic casamino medium (SC; 0.67% yeast nitrogen base, 2% glucose, and 0.5% casamino acids; if necessary, tryptophan, uracil, or adenine was added).

We used mainly W303 strains and its derivative strains. Lists of yeast strains, plasmids, and oligonucleotides are provided in Tables S1, S2, S3, respectively.

For the live-cell imaging of a Hfl1-ymNeongreen (Hfl1-NG), W303 strain with Ade+ *and RAD5* (Y1508), was first created by crossing strain BY20222 (W303, *RAD5*) and Y1489 (W303, Ade+). Then, the strain expressing Hfl1-NG was produced from strain Y1508 by introducing a DNA fragment obtained by PCR using plasmid pFA6a-link-ymNeongreen-SpHis5 (Addgene, #125704) and oligonucleotides #1706, #1707, #1708, and #1709.

To create strain HFL1-3xFlag::HIS3(Y1807); first, a pBSII-3xFlag-TCyc1-HIS3(V165) was created by cloning HIS3 fragment from pRS313 instead of KanMX6 of pBSII-3xFlag-TCyc1-KanMX6 (kindly provided by Dr. H. Yashiroda). A DNA fragment of HFL1-3xFlag-HIS3-HFL1 was obtained by PCR using oligonucleotides #1845, #1846, and using the V165 plasmid as template. The obtained fragment was introduced to W303 to create a strain HFL1-3xFlag::HIS3(Y1807).

HFL1 DNA fragment was obtained by PCR using oligonucleotides #1967 and #1968, cloned into pRS316 (E1055), and subcloned into pRS306 (E1058). To create an *hfl1*(W371A, I375A, D384A, Y387A)-3xFlag::HIS3 strain (Y1916); first, the pRS306-hfl1(D384A Y387A) plasmid (E1059) was created using oligonucleotides #1985 and #1986, using plasmid E1058 as template. Next, pRS306-*HFL1* (D384A, Y387A, W371A, I375A) (E1068) was obtained using oligonucleotides 2114 and 2115 and plasmid E1059. To obtain the hfl1 (D384A, Y387A, W371A, I375A)-3xFlag::HIS3::hfl1 fragment, two PCR fragments were independently obtained using oligonucleotides #2101, #2102 and plasmid E1068 as template and oligonucleotides #2103, #2099 and the genome DNA of strain HFL1-3xFlag::HIS3(Y1807) as template. The two fragments were ligated by Gibson Assembly protocol, amplified using #2101 and #2099, and transformed into W303.

### Freeze fracture EM

Freeze fracture EM analysis was performed as described previously (Tsuji et al., 2017). Briefly, yeast cells sandwiched between a 20-μm–thick copper foil and a flat aluminum disc (Engineering Office M. Wohlwend, Sennwald, Switzerland) were quick-frozen by high-pressure freezing using an HPM 010 high-pressure freezing machine according to the manufacturer’s instructions (Leica Microsystems, Wetzlar, Germany). Frozen specimens were transferred to the cold stage of a Balzers BAF 400 apparatus and fractured at -120° under a vacuum of ~ 1 × 10^−6^ mbar. Freeze-fractured samples were subjected to a three step electron-beam evaporation: C (6 nm) at 90°, Pt/C (2 nm) at 45°, and C (10 nm) as previously described (Tsuji et al., 2017). Thawed replicas were treated with 2.5% SDS in 0.1 M Tris-HCl (pH 8.0) at 60°C overnight, with 0.1% Westase (Takara Bio, Kusatsu, Japan) in McIlvain buffer (37 mM citrate, 126 mM disodium hydrogen phosphate, pH 6.0) containing 10 mM EDTA, 30% fetal calf serum, and a protease inhibitor cocktail for 90 min at 30°C, with 2.5% SDS again in 0.1 M Tris-HCl (pH 8.0) at 60°C overnight. Replicas were observed and photographed with a JEOL JEM-1011 EM (Tokyo, Japan) using a CCD camera (Gatan, Pleasanton, CA, USA).

### Microscopy

FM4-64 staining was performed as described previously (Vida and Emr, 1995); the cells were treated with FM 4-64 just before the temperature shift. For treatment with FM4-64, a 1.5–3 mL culture of cells at early-log phase was grown at 25°C in YPAD medium, followed by centrifugation and suspension in 49 μL of YPAD. To said cells, 1 μL of 2 mM FM4-64 (Invitrogen, T13320) was added at a final concentration of 40 μM and incubated for 20 min at room temperature. The cells were then washed with 1 mL of YPAD and suspended in 1.5–3 mL of YPAD, followed by heat treatment. After the heat treatment, the cells were collected by centrifugation and placed in a heat block until being imaged. Cells were imaged at room temperature using a confocal microscope (LSM700; Carl Zeiss) equipped with a 100× oil objective lens (for images of Figs 1A, 1E, 2A–C, 3A, 6A, and 7A). Images were processed and brightness and contrast were adjusted using Zen software. Alternatively, cells were imaged using a fluorescent microscope (Olympus BX51) equipped with a CMOS camera (ORCA-Fusion, Hamamatsu Photonics), a filter-wheel (Prior, HF110A), and a 100x oil objective lens (UPlan Apo. N/A1.45) (for images of Figs 1D, 4A, and 7C). When observed with this microscopy, cells were washed with filtrated YPD and suspended in YPD before imaging to reduce the background fluorescence from YPD. Images were processed with cellSens software and Photoshop or GIMP. For quantification, cells with invaginated vacuole structures were counted in ≥3 independent experiments.

### Fast Dil staining of vacuoles from heat-stressed yeasts

After growing them at 40.5°C for 2.5 h, cells were harvested by 3,000 rpm for 5 min, washed with spheroplast buffer [1.2M sorbitol, 10 mM K-PO4 (pH7.5)]. and suspended in 3 ml spheroplast buffer containing 1mg/ml zymolyase 100T (Nacalai tesque, 07665-55). Cells were incubated at 40.5°C for 30 min, centrifuged at 3,000 rpm for 5 min, and suspended in 100 μl of lysis buffer (0.2M sorbitol, 10 mM K-PO_4_ (pH7.5)). When cells grown at 25°C were examined, a zymolyase treatment was performed at 30°C for 1 h. Fast Dil (Thermo Fisher D7756) was added at the final concentration of 5 μg mL^-1^ from the stock solution (1 mg mL^-1^ DMSO), and cells were imaged with the Olympus BX51 fluorescent microscopy. Images were processed by deconvolution with cellSens software and GIMP or Photoshop.

### Immunoblotting

The preparation of cell extracts was based on previously described methods (Li et al., 2015; Oku et al., 2017). Briefly, about 1.5 × 10^7^ cells were suspended in 1 mL of ice-cold Buffer A (0.2M NaOH, 0.5%[vol/vol] 2-mercaptoethanol), and one-tenth volume of trichloroacetic acid was added. After 30 min on ice, cells were centrifuged at 20,000 g for 5 min, and the cell pellets were washed twice with 1 mL of cold acetone. Cell pellets were air-dried, suspended in 100 μL of 2x urea buffer (150 mM Tris [pH6.8], 6M urea, and 6% SDS), and glass beads were added. After 5 min of incubation at 37°C, cells were vortexed for 5 min. Then, 100 μL of 2x sample buffer (150 mM Tris [pH6.8], 2% SDS, 100 mM DTT, and bromophenol blue) was added, incubated at 37°C for 5 min, and vortexed again for 5 min. The cell suspension was cooled on ice and centrifuged at 20,000 × g for 1 min just before application and the supernatant was applied to a SDS-PAGE gel.

Western blots were incubated with horseradish peroxidase (HRP)-conjugated anti-Flag antibody (Sigma) or an anti-phosphoglycerate kinase (Life technologies, Frederick MD) followed by horseradish peroxidase (HRP)-conjugated anti-mouse IgG (GE Healthcare, NA931V), and then visualized using a chemiluminescent reagent.

## Data Availability

No datasets were generated or analyzed during the current study.

## Acknowledgements

We thank A. Ishii, A. Koyama, K. Hayashi for their initial observations and N. Shibahara for her technical support. We also thank, Y. Ohsumi, T. Ushimaru, A. Yamamoto, H. Yashiroda, Sakai, Oku, and Y. Jin for materials and advice. We are grateful to S. Yamazaki for discussion. Strain BY20695 was provided by the National Bio-Resource Project of the MEXT, Japan. This work was supported by the Ohsumi Frontier Science Foundation (to Y.K.), Takeda Science Foundation (to Y.K.), Grants-in-Aid for Scientific Research (grant number 20K06620 to Y.K., 19K07265, 22K06818, 22H04654 to T.T.) from the Ministry of Education, Culture, Sports, Science and Technology of Japan (to K.K.) and JST CREST (Grant Number JPMJCR20E3 to T.T.).

